# The anterior insular cortex associates temporally discontiguous stimuli during threat learning

**DOI:** 10.1101/2021.10.19.465048

**Authors:** Ho Namkung, Sedona Lockhart, Josephine de Chabot, Lauren Guttman, Imad Isehak, Hyung-Bae Kwon, Akira Sawa

**Author notes:** Correspondence to: Akira Sawa.

## Abstract

Learning about potential threats in the environment is indispensable for survival. Deficits in threat learning constitute a key dimension of multiple brain disorders, which include posttraumatic stress disorder and anxiety disorder. While human brain imaging studies have highlighted a reliable engagement of the anterior insular cortex (AIC) in threat learning, its precise role remains elusive partly due to the lack of animal studies that can address causality and mechanistic questions. Filling in this gap, the present mouse study proposes a novel AIC-mediated mechanism underlying the association of temporally discontiguous stimuli during threat learning. We identified that activity of AIC layer 5 (L5) pyramidal neurons is required for associating temporally discontiguous stimuli, specifically during a time interval between them. Notably, the AIC is not required for associating temporally contiguous stimuli during threat learning. The AIC not only sends the essential information, via its L5 pyramidal neurons, to the basolateral amygdala (BLA) during the time interval, but also receives from the BLA. We also identified a modulatory role of AIC dopamine D1 receptor (D1R)-mediated dopamine signaling in associating temporally discontiguous stimuli during the time interval.

---

Learning about potential threats in the environment is indispensable for survival (*1*). The brain learns important predictors of threats to optimize future defensive behaviors (*1*). Deficits in threat learning constitute a key dimension of multiple brain disorders, which include posttraumatic stress disorder and anxiety disorder (*2, 3*). Threat learning has been commonly studied in both humans and laboratory animals using fear conditioning paradigms, in which a biologically innocuous stimulus (conditioned stimulus, CS) acquires emotional salience and evolves as a potent predictor of a harmful event (unconditioned stimulus, US) generally through repeated CS-US pairings (*4*). Supported by robust and reproducible physiological/behavioral measures of fear conditioning, the neural substrates that underlie threat learning have been intensively studied in both humans and animals (*5, 6*).

Converging evidence from human brain imaging studies has highlighted that the insular cortex is reliably engaged in threat learning (*6-8*). Nevertheless, different from the amygdala (*9, 10*) and hippocampus (*11*) which are also involved in the process, the precise role of the insular cortex in threat learning remains elusive. This is partly because animal studies to address this important question on the insular cortex are underdeveloped, although animal models are useful in addressing causality and mechanistic questions in principle (*12*). A recent surge of interest in the insular cortex in animals (*13*), following a rich array of human studies (*12*), has started to fill this knowledge gap. However, these efforts have primarily focused on characterizing the posterior insular cortex (PIC) on threat learning (*14-17*), whereas only a few studies have investigated the role of the anterior insular cortex (AIC). Like the amygdala and hippocampus, sub-regions of the insular cortex have distinct roles with each other. The insular cortex is frequently subdivided into posterior and anterior sections: the former with a classical six-layered structure, whereas the latter with a layer IV-lacking structure, called the agranular insular cortex (*12, 13*). While the granular PIC receives heavy sensory inputs across multiple modalities, the agranular AIC further processes this information, interacts with frontal and limbic systems, and supports higher-order cognitive and emotional functions (*12, 13, 18*).

Filling in this fundamental gap, the present study investigates the role of the mouse AIC in threat learning by taking advantage of two complementary paradigms of fear conditioning (FC): delay FC and trace FC. In delay FC, an US is administered to co-terminate with or immediately after a CS, whereas in trace FC a time interval (often called a ‘trace interval’) is introduced between the termination of the CS and the start of the US (*4, 19*). The insertion of a trace interval in trace FC is thought to impose cognitively demanding processes for learning (e.g., holding a ‘trace’ of the first stimulus until the subsequent one occurs), in which brain systems responsible for the higher-order processes may be recruited (*20, 21*). We expect that the trace FC paradigm enables us to investigate the neurobiological basis for associative learning of temporally discontiguous stimuli that we frequently encounter in real-world situations, which may not be captured by the delay FC paradigm.

We first questioned whether AIC activity during conditioning is necessary for threat learning or memory formation. To address this question, we bilaterally infused a mixture of muscimol (GABA_A_ receptor agonist) and baclofen (GABA_B_ receptor agonist) into the AIC before conditioning (*22*) in order to inactivate the AIC in both the delay FC and trace FC paradigms. The inactivation of the AIC did not affect freezing responses to the CS during recall 24 h later in the delay FC paradigm (**Fig. 1, A and B**). In contrast, in the trace FC paradigm, inactivation of the AIC resulted in a significant decrease in freezing responses to the CS during recall (**Fig. 1C**). Given that the infusion of muscimol and baclofen prior to conditioning might also continuously interfere with post-conditioning memory consolidation processes (*23*), we next asked whether the observed failure in threat memory formation in the trace FC paradigm was attributed to a defect in AIC-mediated memory consolidation processes. A bilateral infusion of anisomycin, a protein synthesis inhibitor that blocks memory consolidation, into the AIC (*24*) immediately after conditioning did not affect freezing responses to the CS during recall (**Fig. 1D**). Together, we showed that the AIC is necessary for threat memory formation in the trace FC paradigm, particularly during conditioning, but not during memory consolidation. We next tested the necessity of AIC activity during recall by bilaterally infusing a mixture of muscimol and baclofen into the AIC before recall. We did not observe any significant difference in freezing responses to the CS during recall in both the delay FC and trace FC paradigms (**Fig. 1E and F**). In summary, the AIC is required for threat memory formation only in the trace FC paradigm, but not in delay FC paradigm, with its requirement being during conditioning.

**Fig. 1.**
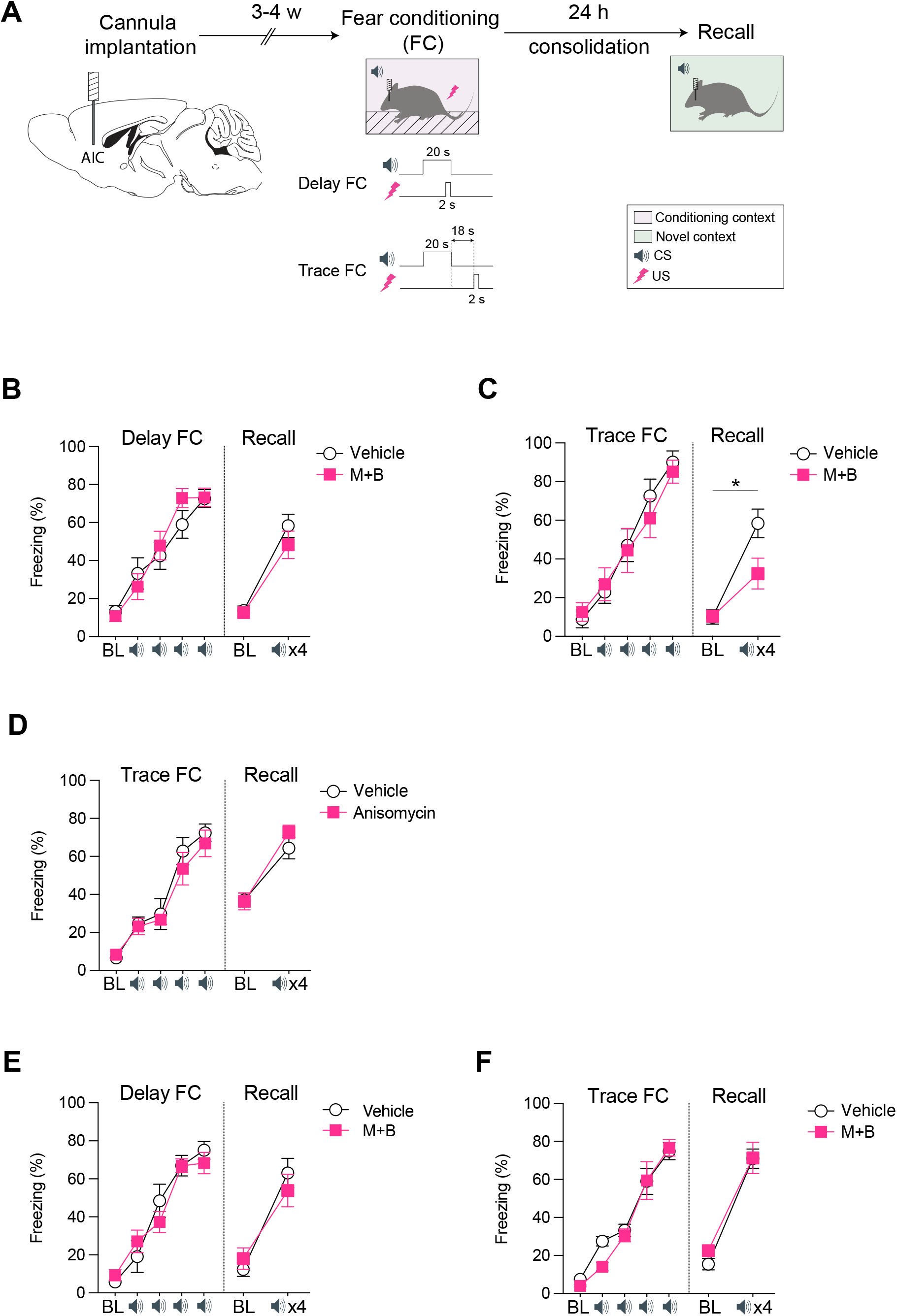
Anterior insular cortex (AIC) activity is required during conditioning for threat memory formation in the trace fear conditioning (FC) paradigm, but not in the delay FC paradigm. **(A)** Schematic of the experimental procedure for pharmacological studies. In delay FC, a 2-s foot-shock (unconditioned stimulus, US) is administered to co-terminate with a 20-s tone (conditioned stimulus, CS), whereas in trace FC an 18-s time interval (often called a ‘trace interval’) is introduced between the termination of the 20-s CS and the start of the 2-s US. **(B)** Pharmacological inactivation of the AIC during conditioning, which was realized by bilaterally infusing a mixture of muscimol and baclofen (M+B) into the AIC 30 min before conditioning, did not affect freezing responses to the CS, during recall 24 h later in the delay FC paradigm. The baseline (BL) response indicates mean percentage freezing during the initial 2-min BL period prior to the first CS presentation. Cued fear recall performance was measured by averaging freezing responses to the 4 CS presentations in a novel context. Delay FC (*N*_Vehicle_=12, *N*_M+B_=12; two-way repeated measures (RM) ANOVA with the Greenhouse-Geisser correction, Drug x CS, *F*_3.027,66.584_=1.455, *p*=0.235). Recall (*N*_Vehicle_=12, *N*_M+B_=12; two-way RM ANOVA, Drug x CS, *F*_1,22_=0.703, *p*=0.411). **(C)** Pharmacological inactivation of the AIC during conditioning resulted in significant decrease in freezing responses to the CS during recall in the trace FC paradigm. Trace FC (*N*_Vehicle_=11, *N*_M+B_=13; two-way RM ANOVA with the Greenhouse-Geisser correction, Drug x CS, *F*_2.946,64.802_=0.596, *p*=0.617). Recall (*N*_Vehicle_=11, *N*_M+B_=13; two-way RM ANOVA, Drug x CS, *F*_1,22_=7.888, *p*=0.01). **(D)** Bilateral infusion of anisomycin into the AIC immediately after conditioning did not affect freezing responses to the CS during recall in the trace FC paradigm. Trace FC (*N*_Vehicle_=9, *N*_Anisomycin_=10; two-way RM ANOVA with the Greenhouse-Geisser correction, Drug x CS, *F*_2.230,37.905_=0.371, *p*=0.715). Recall (*N*_Vehicle_=9, *N*_Anisomycin_=10; two-way RM ANOVA, Drug x CS, *F*_1,17_=3.142, *p*=0.094). **(E)** Pharmacological inactivation of the AIC during recall did not affect freezing responses to the CS in the delay FC paradigm. Delay FC (*N*_Vehicle_=10, *N*_M+B_=12; two-way RM ANOVA with the Greenhouse-Geisser correction, Drug x CS, *F*_2.787,55.744_=1.084, *p*=0.360). Recall (*N*_Vehicle_=10, *N*_M+B_=12; two-way RM ANOVA, Drug x CS, *F*_1,20_=1.646, *p*=0.214). **(F)** Pharmacological inactivation of the AIC during recall did not affect freezing responses to the CS in the trace FC paradigm. Trace FC (*N*_Vehicle_=10, *N*_M+B_=10; two-way RM ANOVA with the Greenhouse-Geisser correction, Drug x CS, *F*_1.803,32.449_ =0.841, *p*=0.430). Recall (*N*_Vehicle_=10, *N*_M+B_=10; two-way RM ANOVA, Drug x CS, *F*_1,18_=0.355, *p*=0.559). \**p* < 0.05. Graph expressed as mean ± SEM.

We therefore asked in which specific time window during conditioning the AIC is required for threat memory formation in the trace FC paradigm. To address this question, we employed time window-specific optogenetic inhibition of the AIC, which enables us to dissect its necessity at a high temporal resolution that cannot be achieved by pharmacological manipulation. We inhibited AIC excitatory neurons selectively during the CS, trace interval (TI), or US, respectively, as well as during the entire period of each trial, by using enhanced halorhodopsin (eNpHR3.0) expressed under the CaMKIIa promoter (**Fig. 2A**). Selective optogenetic inhibition of the AIC excitatory neurons during the TI prevented threat memory formation, whereas optogenetic inhibition during CS or US did not (**Fig. 2B**). Meanwhile, inhibition during the entire period also prevented threat memory formation (**Fig. 2B**). Together, for threat memory formation in the trace FC paradigm, AIC excitatory neuronal activities selectively during the TI are crucial. In contrast, optogenetic inhibition of the PIC excitatory neurons did not disrupt threat memory formation, indicating the distinct role of the AIC from the PIC (**Fig. S1A**).

**Fig. 2.**
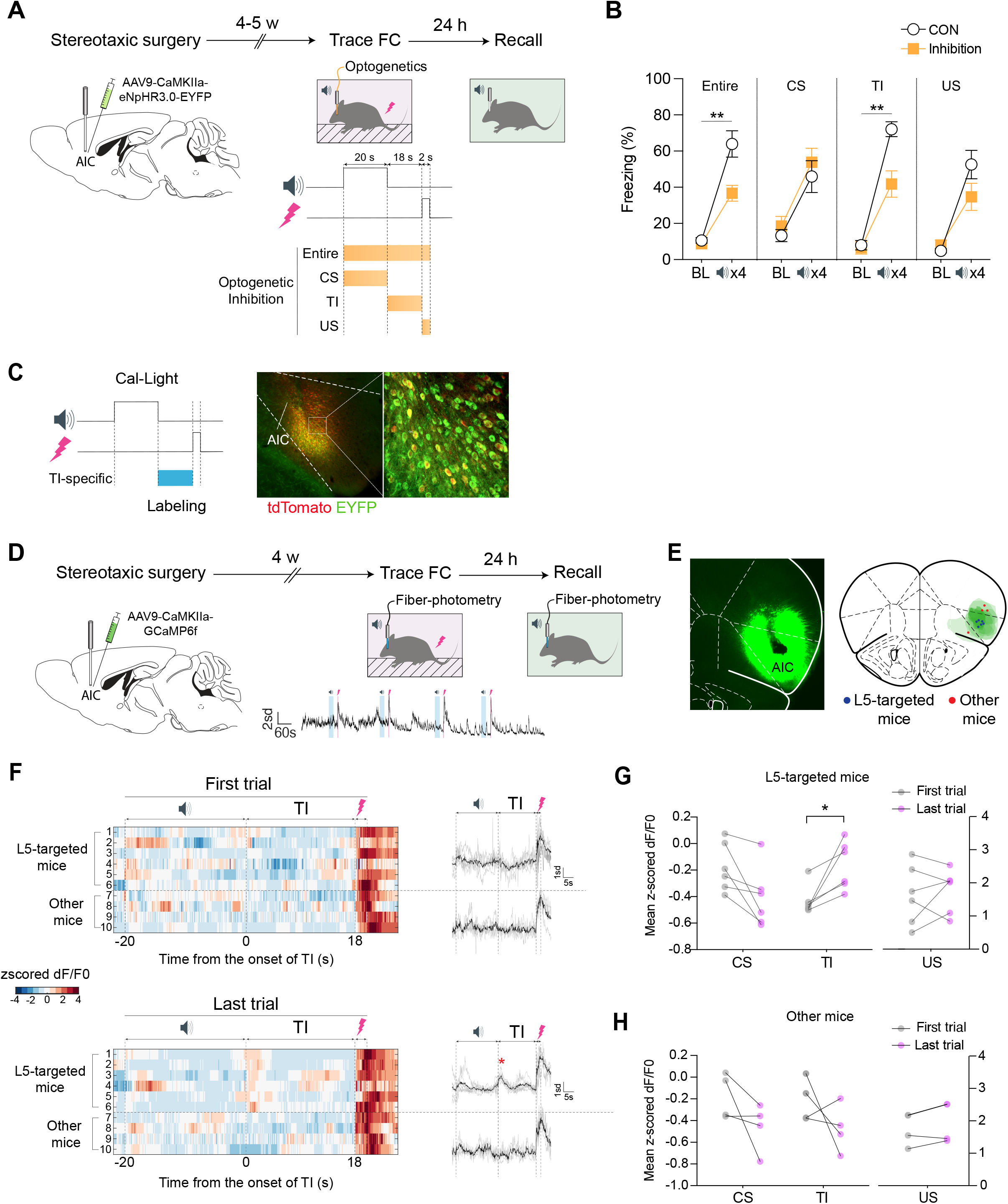
AIC excitatory neuronal activity during the trace interval (TI) is required for threat memory formation in the trace FC paradigm. **(A)** Schematic of the experimental procedure for time window-specific optogenetic inhibition of AIC excitatory neurons. **(B)** Optogenetic inhibition of AIC excitatory neurons during the 18-s TI as well as during the entire period of each trial, but not during the 20-s CS or 2-s US, prevented threat memory formation in the trace FC paradigm. Entire (*N*_CON_=13, *N*_Inhibition_=12; two-way RM ANOVA, Inhibition x CS, *F*_1,23_=9.483, *p*=0.005). CS (*N*_CON_=11, *N*_Inhibition_=13; two-way RM ANOVA, Inhibition x CS, *F*_1,22_=0.268, *p*=0.610). TI (*N*_CON_=9, *N*_Inhibition_=10; two-way RM ANOVA, Inhibition x CS, *F*_1,17_=12.281, *p*=0.003). US (*N*_CON_=12, *N*_Inhibition_=14; two-way RM ANOVA, Inhibition x CS, *F*_1,24_=4.254, *p*=0.06). **(C)** The AIC neurons activated specifically during the TI were shown to be enriched in the AIC layer 5 (L5) by introducing a light-inducible activity-dependent gene expression system, named Cal-Light. **(D)** Schematic of the experimental procedure for *in vivo* fiber photometry to record AIC excitatory neuronal dynamics. **(E)** Histological assessment of the implantation sites of fiber optic cannulas. Subjects were clustered into two groups based on the implantation sites of fiber optic cannulas: the group of L5-targeted mice and the other group implanted outside L5. **(F)** The population activity of AIC excitatory neurons was compared between the first and last trials. The group of L5-targeted mice showed transient AIC neuronal activity (indicated as a red asterisk) time-locked to the onset of the TI on the last trial. **(G)** The group of L5-targeted mice showed a significant increase in average neuronal activity during the TI, but not during the CS or US, when the first and last trials were compared. CS (*N*_First_=6, *N*_Last_=6; two-tailed paired student’s *t*-test, *t*_5_=1.935, *p*=0.082). TI (*N*_First_=6, *N*_Last_=6; two-tailed paired student’s *t*-test, *t*_5_=-3.430, *p*=0.019). US (*N*_First_=6, *N*_Last_=6; two-tailed paired student’s *t*-test, *t*_5_=-0.476, *p*=0.654). **(H)** The neuronal activity-associated signature during the TI was not observed in non L5-targed mice. CS (*N*_First_=4, *N*_Last_=4; two-tailed paired student’s *t*-test, *t*_3_=1.663, *p*=0.195). TI (*N*_First_=4, *N*_Last_=4; two-tailed paired student’s *t*-test, *t*_3_=0.082, *p*=0.940). US (*N*_First_=4, *N*_Last_=4; two-tailed paired student’s *t*-test, *t*_3_=-1.998, *p*=0.140). \**p* < 0.05, \*\**p* < 0.01. Graph expressed as mean ± SEM.

We next questioned whether a specific AIC neuronal population is engaged for threat memory formation selectively during the TI in the trace FC paradigm. To address this question, we labeled neurons that were activated in the AIC only during the TI by using Cal-Light, a light-inducible activity-dependent gene expression system (*25*). This system can drive expression of the reporter EGFP only in active neurons with high spatiotemporal resolution in the presence of blue light. Accordingly, we identified that the neurons activated during the TI are enriched in the AIC layer 5 (L5) (**Fig. 2C**). We therefore investigated the underlying dynamics of the AIC L5 pyramidal neurons throughout the course of trace FC, with a particular focus on the TI. We conducted *in vivo* fiber photometry to record AIC excitatory neuronal dynamics by expressing the fluorescent calcium indicator GCaMP6f under the CaMKIIa promoter (**Fig. 2D**). We then clustered subjects into two groups based on the implantation sites of fiber optic cannulas: the group of L5-targeted subjects and the other group implanted off target (**Fig. 2E**). We identified that the group of L5-targeted mice, but not the other, showed transient AIC neuronal activity time-locked to the onset of the TI on the last trial (**Fig. 2F**). In line with this observation, there was a significant increase in average neuronal activity during the TI, but not during the CS or US, when we compared between the first and last trials (**Fig. 2G**). However, this neuronal activity-associated signature during the TI was not observed in non-L5 neurons (**Fig. 2H**).

Based on these observations in the AIC, we next investigated a circuitry mechanism involving long-range projection pathways by which the AIC may play an essential role in threat memory formation in the trace FC paradigm. We hypothesized that the TI-specific activity of the AIC L5 pyramidal neurons might jointly contribute to threat memory formation with cortical and subcortical brain regions that receive dense projections from the AIC L5 pyramidal neurons and are also involved in this paradigm (*26, 27*). Accordingly, we focused on the basolateral amygdala (BLA), dorsomedial prefrontal cortex (dmPFC), and hippocampus (HP). We observed strong axonal terminals within the BLA and dmPFC, but not the HP, by anterograde tracing (**Fig. 3A**). We next tested whether the AIC-to-BLA (AIC→BLA) and/or the AIC-to-dmPFC (AIC→dmPFC) excitatory inputs, arising from AIC L5 pyramidal neurons, were functionally required for threat memory formation selectively during the TI. To address this question, we used projection-specific optogenetic inhibition by expressing a mosquito-derived rhodopsin (eOPN3) in the AIC of Retinol Binding Protein 4 (RBP4)-Cre mice in a Cre-dependent manner, and photo-inhibited their axonal terminals in either the BLA or dmPFC selectively during TI. We observed that TI-specific silencing of the AIC→BLA pathway, but not the AIC→dmPFC pathway, prevented threat fear memory formation, indicating the required activity of the AIC L5 pyramidal neurons projecting to the BLA (**Fig. 3B**).

**Fig. 3.**
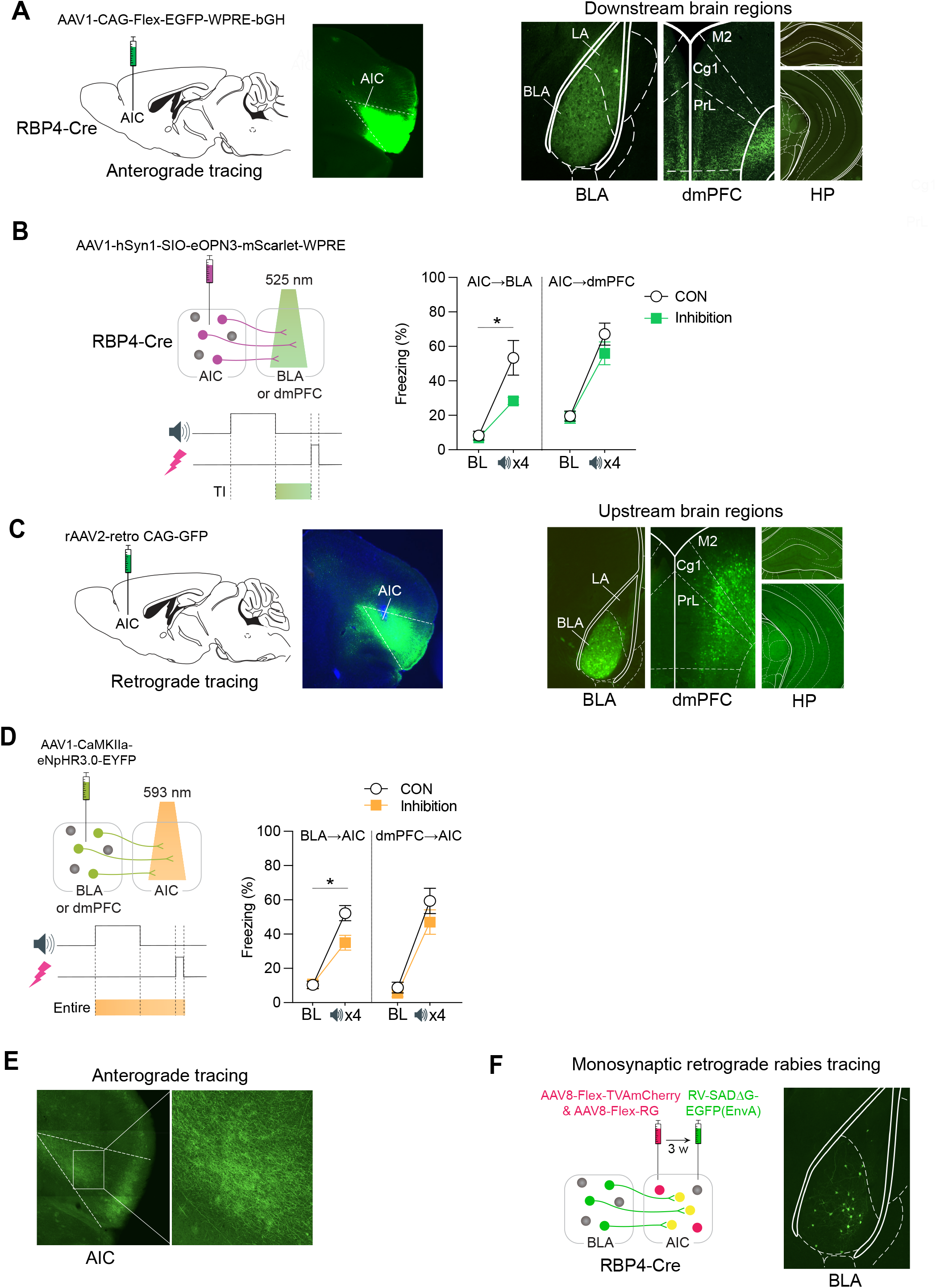
The reciprocal interactions between the AIC and the basolateral amygdala (BLA) are required for threat memory formation in the trace FC. **(A)** Anterograde tracing of axonal targets of RBP4-positive neurons in the AIC, with a particular focus on three downstream brain regions: the BLA, dorsomedial prefrontal cortex (dmPFC), and hippocampus (HP). **(B)** TI-specific silencing of the AIC-to-BLA (AIC→BLA) excitatory inputs, but not the AIC-to-dmPFC (AIC→dmPFC) excitatory inputs, which arise from the AIC L5 pyramidal neurons, prevented threat fear memory formation in the trace FC paradigm. AIC→BLA (*N*_CON_=10, *N*_Inhibition_=10; two-way RM ANOVA, Inhibition x CS, *F*_1,18_=7.144, *p*=0.016). AIC→dmPFC (*N*_CON_=12, *N*_Inhibition_=12; two-way RM ANOVA, Inhibition x CS, *F*_1,22_=1.631, *p*=0.215). **(C)** Expression of retrogradely-transported GFP into the AIC identified enriched AIC-projecting neurons within the BLA and dmPFC, but not the HP. **(D)** Optogenetic silencing of the BLA→AIC excitatory pathway, but not the dmPFC→AIC excitatory pathway, during the entire period of each trial prevented threat fear memory formation in the trace FC paradigm. BLA→AIC (*N*_CON_=8, *N*_Inhibition_=8; two-way RM ANOVA, Inhibition x CS, *F*_1,14_=8.094, *p*=0.013). dmPFC→AIC (*N*_CON_=10, *N*_Inhibition_=10; two-way RM ANOVA, Inhibition x CS, *F*_1,18_=0.855, *p*=0.367). **(E)** Anterograde labeling of BLA axons identified enriched terminals in the AIC deep layers (L5 and L6). **(F)** Cell type-specific, rabies virus-based retrograde tracing of RBP4-positive neurons in the AIC validated the enrichment of the BLA neurons that directly project to the AIC L5 pyramidal neurons. \**p* < 0.05. Graph expressed as mean ± SEM.

We next mapped afferent inputs to the AIC by expressing retrogradely-transported GFP into the AIC, and identified enriched AIC-projecting neurons within the BLA and dmPFC, but not the HP (**Fig. 3C**). To address the possible necessity of BLA→AIC and/or dmPFC→AIC excitatory inputs for threat memory formation in the trace FC paradigm, we utilized projection-specific optogenetic inhibition. We expressed eNpHR3.0 in either the BLA or dmPFC under the CaMKIIa promoter and then photo-inhibited the axonal terminals in the AIC during the entire period of each trial. We found that optogenetic silencing of the BLA→AIC pathway, but not the dmPFC→AIC pathway, prevented threat fear memory formation, indicating the requirement of BLA excitatory inputs to the AIC (**Fig. 3D**). Anterograde labeling of BLA axons identified enriched terminals in the AIC deep layers (L5 and L6) (**Fig. 3E**). Furthermore, cell type-specific, rabies virus-based retrograde tracing of RBP4-positive neurons in the AIC validated the enrichment of the BLA neurons that directly project to the AIC L5 pyramidal neurons (**Fig. 3F**). Altogether, a reciprocal interaction between the AIC and BLA is necessary for the association of the CS and US across the TI in the trace FC paradigm.

What can be a potential modulator of the AIC-mediated threat memory formation during the TI in communication with the BLA in the trace FC paradigm? Dopaminergic modulation of the prefrontal cortex (PFC) has been highlighted in higher-order cognitive processes (*28*). Dopamine D1 receptor (D1R) stimulation in the PFC is known to produce an ‘inverted-U’ dose-response: either too much or too little D1R stimulation impairs working memory (*28, 29*). Accordingly, we hypothesized that dopaminergic modulation via the D1R might underlie the aforementioned function of the AIC in threat memory formation. We observed that infusion of the D1R antagonist SCH23390 in the AIC before conditioning reduced fear responses to the CS during both conditioning and recall, but the D2R antagonist sulpiride did not (**Fig. 4A**). These data support a modulatory role of D1R-mediated dopamine signaling in the AIC-mediated threat learning and memory formation. We next validated whether dopamine signaling is necessary during the TI by using cell type-specific optogenetic inhibition. We observed that the TI-specific inhibition of D1R-positive neurons in the AIC significantly weakened threat memory formation in the trace FC paradigm (**Fig. 4B**). We next addressed dopamine dynamics in the AIC throughout the course of trace FC by employing *in vivo* fiber photometry with a dopamine indicator dLight1.1, which represents the conformational changes of dopamine receptors in response to dopamine binding (*30*). We observed a reduction in dLight1.1 signals in the AIC during the TI, but not during the CS or US, when we compared between the first and last trials (**Fig. 4, C and D**). Together, these results indicate that the modulation of AIC dopamine signaling during the TI has a significant role, likely via D1R-positive neurons, in threat memory formation in the trace FC paradigm.

**Fig. 4.**
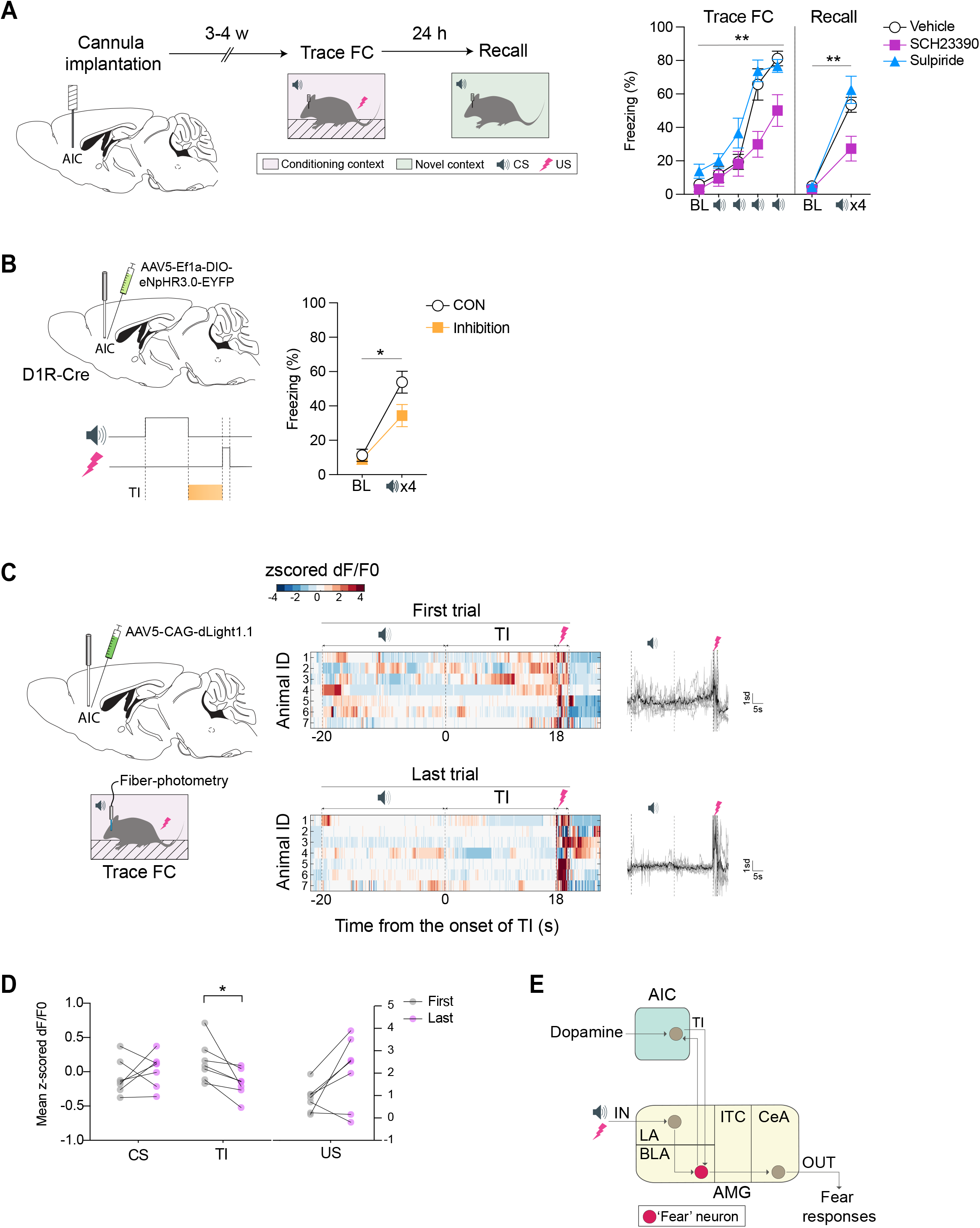
Dopamine D1 receptor (D1R)-mediated DA signaling in the AIC, specifically during the TI, is necessary for threat memory formation in the trace FC paradigm. **(A)** Infusion of the D1R antagonist SCH23390 in the AIC 30 min before conditioning reduced fear responses to the CS during both conditioning and recall, but the D2R antagonist sulpiride did not. Trace FC (*N*_CON_=9, *N*_SCH23390_=8, *N*_Sulpiride_=9; two-way RM ANOVA, Drug x CS, *F*_8,92_=3.359, *p*=0.002; *post hoc* Bonferroni test, *p*_CON vs. SCH23390_=0.046, *p*_CON vs. Sulpiride_=0.563). Recall (*N*_CON_=9, *N*_SCH23390_=8, *N*_Sulpiride_=9; CSx4, two-way RM ANOVA, Drug x CS, *F*_2,23_ =6.810, *p*=0.005; *post hoc* Bonferroni test, *p*_CON vs. SCH23390_=0.038, *p*_CON vs. Sulpiride_=1.0). **(B)** TI-specific inhibition of D1R-positive neurons in the AIC significantly weakened threat memory formation in the trace FC paradigm. TI (*N*_CON_=13, *N*_Inhibition_=13; two-way RM ANOVA, Inhibition x CS, *F*_1,24_=4.468, *p*=0.045). **(C)** Dopamine dynamics in the AIC, throughout the course of trace FC, were examined by employing *in vivo* fiber photometry with a dopamine indicator dLight1.1. **(D)** The dLight1.1 signals in the AIC were significantly reduced during the TI, but not during the CS or US, when we compared between the first and last trials. CS (*N*_First_=7, *N*_Last_=7; two-tailed paired student’s *t*-test, *t*_6_=-0.841, *p*=0.432). TI (*N*_First_=7, *N*_Last_=7; two-tailed paired student’s *t*-test, *t*_6_=2.810, *p*=0.031). US (*N*_First_=7, *N*_Last_=7; two-tailed paired student’s *t*-test, *t*_6_=-2.286, *p*=0.062). **(E)** A novel AIC-mediated mechanism underlying the association of temporally discontinuous stimuli during threat learning. LA: lateral amygdala; ITC: intercalated amygdala; CeA: central amygdala; AMG: amygdala; IN: input; OUT: output. \**p* < 0.05, \*\**p* < 0.01. Graphs expressed as mean ± SEM.

The present study proposes a novel AIC-mediated mechanism underlying the association of temporally discontiguous stimuli during threat learning (**Fig. 4E**). The requirement of the AIC L5 pyramidal neurons for threat memory formation only in the trace FC paradigm, but not in the delay FC paradigm, suggests an essential role of the AIC in mediating some higher-order process specifically during the TI. In contrast, the PIC excitatory neurons were not required for this process, indicating that this AIC-mediated role may represent a higher-order function unique to the AIC, which is not mediated by the PIC.

We also found that reciprocal communications between the AIC and BLA are required for threat memory formation in the trace FC paradigm. While the BLA is well-known to be essential for associative learning of the CS and US in delay FC, it remains elusive how in trace FC the BLA associates the CS and US that are temporally separated by the TI. Our study proposes a novel role of the AIC L5 pyramidal neurons in the integration of temporally discontiguous information through the reciprocal communication with the BLA. However, future investigation needs to address how these two brain regions communicate particularly in the temporal sequences.

We also identified a modulatory role of D1R-mediated dopamine signaling in the AIC for threat learning and memory formation in the trace FC paradigm. In addition, we observed a significant reduction in dopamine dynamics selectively during the TI in the AIC. According to the dopamine “gating” theory of working memory, the cortical dopamine level can gate the update of memory contents by operating in two modes: an updating (gate-open) mode, which allows new information to enter working memory, and a maintenance (gate-closed) mode, which prevents distracting information from interfering with the current contents of working memory (*31, 32*). The reduction in dopamine dynamics during the TI in the AIC may temporarily change the gating mode of the AIC so that the process of associating temporally discontiguous stimuli (CS and US) during the TI is protected from distracting noises. It is also unknown what drives the TI-specific change in dopamine dynamics in the AIC. Further investigation for this potential gating mechanism is warranted.

Through human brain imaging studies, a significant involvement of the AIC has been reproducibly underscored in many neurological and psychiatric disorders, which include not only posttraumatic stress disorder and anxiety disorder but also schizophrenia (*12, 33*). The AIC is also well known as a key hub of the salience network, which is frequently discussed in the pathophysiological contexts (*12, 34*). It is conceivable that AIC-mediated bridging of temporal discontiguity may underlie the formation of salient associations. Therefore, the novel function of the AIC reported here may participate in the mechanism for aberrant salience that underlie a wide range of neurological and psychiatric disorders.

## Acknowledgments

We thank Drs. Patricia Janak, Seung-Chan Lee, and John Issa for insightful comments on the manuscript. We thank Dr. Solange Brown for sharing RBP4-Cre mice. We thank Victoria Stepanyants and Brian Lee for a pilot electrophysiological investigation.

## Author contributions

H.N. formulated the conceptual framework, designed and conducted experiments, and wrote the manuscript. S.L., J.C., L.G., I.I. conducted experiments. H-B.K. assisted conceptual framework formulation and experimental design in neurobiological study. A.S. formulated the conceptual framework, supervised the overall project, and wrote the manuscript

## Data and materials availability

All major data generated or analyzed are included in the article. Data and materials related to this study are available from the corresponding author on reasonable request.

