## Supplementary materials for "The anterior insular cortex associates temporally discontiguous stimuli during threat learning"

#### Materials and Methods

##### Subjects

Adult male mice at 9-18 weeks old were used in this study. Wild-type C57BL/6J mice (Jackson Laboratories) were used for pharmacological inactivation, optogenetic inhibition, fiber photometry, antero-/retro-grade tracing, and light-inducible activity-dependent gene expression. RBP4-Cre mice (*35*) on the C57BL/6J background, which are shown to express Cre in layer 5 pyramidal neurons (*36*), were used for cell type-specific optogenetic terminal inhibition of RBP4-positive neurons and antero-/retro-grade viral tracings. Drd1-Cre mice (GENSAT BAC transgenic EY262) (*37*) on the C57BL/6J background were used for cell type-specific optogenetic inhibition of D1R-positive neurons. All experimental procedures were in accordance with NIH guidelines and approved by the Johns Hopkins Animal Care and Use Committee.

##### Viral constructs and stereotaxic surgeries

For fiber photometry of AIC excitatory neuronal population dynamics, AAV9-CaMKII-GCaMP6f-WPRE-SV40 (0.5 ul, titer: 2.3 x $10^{13}$ GC/mL, 100834-AAV9, Addgene) was injected unilaterally into the AIC of wild-type mice at the following coordinate from Bregma: AP +2.34 mm, ML -2.3 mm, DV -2.7 mm. A fiber optic cannula (MFC_400-430-0.48_5mm, Doric lenses) was implanted into the same site and secured to the skull with acrylic glue. For fiber photometry of AIC DA dynamics, the same AIC coordinate was targeted with AAV5-CAG-dLight1.1 (0.5 ul, titer: 8.1 x $10^{12}$ GC/mL, 111067-AAV5, Addgene). A fiber optic cannula was implanted into the same site and secured to the skull with acrylic glue. For optogenetic inhibition of AIC excitatory neurons, AAV9-CaMKIIa- eNpHR3.0-EYFP (0.5 ul, titer: 9.66 x $10^{12}$ GC/mL, 26971-

AAV9, Addgene) or as a control AAV5-CaMKIIa-EYFP (0.5 ul, titer: 2.5 x $10^{13}$ GC/mL, 50469-AAV5, Addgene) was injected bilaterally into the AIC at the following coordinates from Bregma: AP +2.34 mm, ML ±2.3 mm, DV -2.7 mm. The fiber optic cannulas were implanted 0.5 mm above the injection sites and secured to the skull with acrylic glue. For optogenetic inhibition of RBP4-positive AIC neurons that project to either the BLA or dmPFC, AAV1-hSyn1-SIO-eOPN3-mScarlet-WPRE (0.5 ul, titer: 1.0 x $10^{13}$ GC/mL, 125713-AAV1, Addgene) or as a control AAV5-Ef1a-DIO-EYFP (0.5 ul, titer: 3.3 x $10^{13}$ GC/mL, 27056-AAV5, Addgene) was injected bilaterally into the AIC of RBP4-Cre mice at the same AIC coordinates. For inhibition of axonal terminals in either the BLA or dmPFC, the fiber optic cannulas were implanted bilaterally at the following coordinates from Bregma and secured to the skull with acrylic glue: BLA (AP -1.22 mm, ML ±3.27 mm, DV-4.2 mm) and dmPFC (AP +1.94 mm, ML ±0.5 mm, DV-1.0 mm). For optogenetic inhibition of either BLA or dmPFC excitatory neurons that project to the AIC, AAV1-CaMKIIa- eNpHR3.0-EYFP (0.5 ul, titer: 9.66 x $10^{12}$ GC/mL, 26971-AAV1, Addgene) or as a control AAV5-CaMKIIa-EYFP (0.5 ul, titer: 2.5 x $10^{13}$ GC/mL, 50469-AAV5, Addgene) was injected bilaterally into either the BLA or dmPFC at the following coordinates from Bregma: BLA (AP -1.22 mm, ML ±3.27 mm, DV-4.7 mm) and dmPFC (AP +1.94 mm, ML ±0.3 mm, DV-1.5 mm). For inhibition of axonal terminals in the AIC, the fiber optic cannulas were implanted bilaterally into the AIC at the following coordinates from Bregma and secured to the skull with acrylic glue: AP +2.34 mm, ML ±2.3 mm, DV -2.2 mm. For optogenetic inhibition of D1R-positive AIC neurons, AAV5-Ef1a-DIO-eNpHR3.0-EYFP (0.5 ul, titer: 2.5 x $10^{13}$ GC/mL, 26966-AAV5, Addgene) or as a control AAV5-Ef1a-DIO-EYFP (0.5 ul, titer: 3.3 x $10^{13}$ GC/mL, 27056-AAV5, Addgene) was injected bilaterally into the AIC of D1R-Cre mice at the following coordinates from Bregma: AP +2.34 mm, ML ±2.3 mm, DV -2.7 mm.

The fiber optic cannulas were implanted bilaterally 0.5 mm above the injection sites and secured to the skull with acrylic glue. For retrograde tracings of afferent inputs to the AIC, rAAV2-retro CAG-GFP (0.5 ul, titer: 1.2 x $10^{13}$ GC/mL, 37825-AAVrg, Addgene) was injected unilaterally into the AIC at the following coordinate from Bregma: AP +2.34 mm, ML -2.3 mm, DV -2.7 mm. For monosynaptic retrograde rabies tracings of RBP4-positive AIC neurons, mice were first unilaterally injected with 0.5 ul of a 6:1 (RG:TV) mixture of helper viruses (AAV8-Flex-RG, titer: 4.6 x $10^{12}$ GC/mL, UNC Vector core; AAV8-Flex-TVAmCherry, titer: 5.4 x $10^{12}$ GC/mL, UNC Vector core) into the AIC of RBP4-Cre mice at the following coordinate from Bregma: AP +2.34 mm, ML -2.3 mm, DV -2.7 mm. Three weeks later, 0.5 ul of RV-SADΔG-EGFP (EnvA) (titer: 2.4 x $10^{8}$ TU/mL, Salk Viral Vector Core) was injected at the same coordinate, and the mice were perfused for histological analysis 7-10 days after rabies virus injections. For any viral injections, mice were deeply anesthetized with an intraperitoneal injection of Avertin (0.4 mg/g) and placed in a stereotaxic apparatus (David Kopf Instruments). The head was leveled using Bregma and Lambda reference points and a craniotomy was performed. Viruses were injected using a syringe (Hamilton, 80383), the syringe was left in place for 5 minutes after each injection, and then it was slowly withdrawn. The skin was sutured and closed. Mice were recovered from anesthesia on a heating pad, and then were returned to their home cage. Animals were allowed at least 3 weeks of recovery time after the surgery, except for some tracing experiments.

Drug infusions

For drug infusions, mice were implanted bilaterally with 26 gauge guide cannulas (8IC315GSPCXC, PlasticsOne) into the AIC at the following coordinates from the Bregma and secured to the skull with acrylic glue: AP +2.34 mm, ML ±2.3 mm, DV -2.7 mm. The intracranial microinjection was conducted using a microsyringe pump controller MICRO-4 (World Precision Instruments) connected with a Hamilton microsyringe (Plastic Ones, HSYR-1) through a tube (Plastics Ones, C230C/SPC) with the flow rate 0.15 uL/min. For pharmacological inactivation of the AIC, a mixture [(0.5 ug + 0.5 ug) / 0.3 ul / side] of Muscimol (028945, Tocris Bioscience) + Baclofen (0417, Tocris Bioscience) or as a control saline was infused bilaterally into the AIC 30 minutes before FC or before recall, depending on the time window of interest. For blocking post-conditioning memory consolidation mediated by the AIC, anisomycin (62.5 ug / 0.3 ul / side in 1 M HCl and 100 mM PBS, pH7.2, A9789-25MG, SIGMA-ALDRUICH) or as a control saline was infused bilaterally into the AIC right after trace FC. For testing the necessity of D1R- or D2R-mediated DA signaling in threat memory formation of trace FC, SCH23390 (1 ug / 0.3 ul / side, CAS 125941-87-9, SIGMA-ALDRUICH), Supliride (1 ug / 0.3 ul / side, S8010-25G, SIGMA-ALDRUICH), or Saline as a control was infused bilaterally into the AIC 30 minutes before trace FC.

Fear conditioning

Delay and trace fear conditioning (FC) paradigms were employed in a comparative manner. Both paradigms were composed of three experimental phases (habituation, fear conditioning, and recall), with each phase being studied on three consecutive days in large sound-proof isolation chambers (Med associates Inc.). On day 1, mice underwent the habituation phase for 10 minutes in a conditioning context. On day 2 (24 h after habituation), in the conditioning context, mice were subject to associate a pure tone (conditioned stimulus, CS; 3 KHz, 70 dB, 20 s) with a foot-shock (unconditioned stimulus, US; 2 s, 0.5 mA) through 4 repeated CS-US pairings, with each paring being separated by a variable interval (160 -200 s). Delay and trace FC differed only in the time interval between the CS and US: the 2-s US was administered to co-terminate with the 20-s CS in delay FC, whereas in trace FC an 18-s time interval (i.e., trace interval) is inserted between the termination of the 20-s CS and the start of the 2-s US. On day 3 (24 h after fear conditioning), cued fear recall performance was measured by recording and averaging freezing responses to the 4 CS presentations in a newly-constructed context (i.e., a novel context) that is distinct from the conditioning context, with each CS being presented at a variable interval (160–200 seconds). Cued fear recall performance was measured by averaging freezing responses to the 4 CS presentations in a novel context. A baseline (BL) response was measured by mean percentage freezing during the initial 2-min BL period prior to the first CS presentation.

Fiber photometry

Fiber photometry was employed to monitor population $ca^{2+}$-sensitive fluorescence or DA dynamics from genetically defined cell types. Two LEDs (465 nm for GCaMP6 excitation and 405 nm for isosbestic wavelength excitation, CLED_465 and CLED_405, Doric lenses) were synchronized using a TTL pulse generator (OTPG4, Doric lenses) to have them work in alternative excitations at 10 Hz. Excitation light was coupled into a low auto-fluorescence 400 um patch cord (0.48 NA, 1 m cable length, Doric lenses) whose end was connected to an fiber optic cannula/ferrule (MFC_400-430-0.48_5mm/ SLEEVE_ZR_1.25-BK, Doric lenses) implanted into the AIC. The LED powers for both excitation wavelengths were measured with a power meter (PM100D, Thorlabs) and set to be 50-80 uW at the end of the patch cord. Emission light from GCaMP6 was collected through the same fiber using a sCMOS camera (Zyla 4.2, Andor), which was synchronized with two LEDs using the same TTL pulse generator to capture signals at 20 Hz and controlled using the software, named Micro-manager. To calculate ΔF/F, a least-squares linear fit was applied to the 405 nm signal to align it to the 465 nm signal, which produces a fitted 405 nm signal used to normalize the 465 nm as follows: ΔF/F = (465 nm signal − fitted 405 nm signal)/fitted 405 nm signal.

Optogenetics

Optogenetics was introduced to manipulate neural activity of genetically defined cell types in the AIC or AIC-connected pathways. For optogenetic inhibition, bilaterally implanted fiber optic cannulas/ferrules (MFC_200-245-0.53_5mm_ZF1.25_FLT/SLEEVE_ZR_1.25-BK, Doric lenses) were connected with the bifurcated ends of a branching fiber-optic patch cord whose another end was connected to a fiber-optic rotary joint (FRJ_1x1_FC-FC, Doric lenses) to allow free movement. Another patch cord was connected from the rotary joint to either a 612 nm laser (Ce:YAG & Blue Fiber Light Source/CF-Custom Filter FF01-612/69, Doric lenses) or a 549 nm laser (Ce:YAG Fiber Light Source/CF-Custom Filter FF01-549/15, Doric lenses) for continuous stimulation of eNpHR3.0 or eOPN3, respectively. The laser power for both excitation wavelengths was measured with a power meter (PM100D, Thorlabs) and set to be 10 mW at the tip. Laser power was measured before and after every FC session to ensure that all equipment was functioning properly. Laser output was synchronized by a TTL pulse generator (OTPG4, Doric lenses) and controlled by the Doric Neuroscience Studio software (Doric lenses).

Histology

Mice were euthanized and transcardially perfused with PBS, immediately followed by perfusion with 4% paraformaldehyde (PFA). Brains were post-fixed overnight in PFA at 4°C and coronally sectioned with a cryostat (Leica) at 50 μm. Brain cryosections were incubated for 16 h at 4°C with primary antibody: anti-GFP (1:500; 3h9-20, Proteintech) or anti-RFP (1:500; 5f8-20, Proteintech) antibodies in 10 % normal goat serum (NGS) in PBST (PBS, 1% BSA, and 0.5% Triton X-100). The sections were then washed and incubated for 2 hours at room temperature with either the secondary Alexa 488 goat anti-rat (1:400; Invitrogen, A11006) antibody or Alexa 568 goat anti-rat (1:400, Invitrogen, A11077) antibody in 1% NGS/PBST. Stained sections were washed and DAPI-stained (Invitrogen). All sections were mounted and imaged on a Zeiss AX10 Observer D1 or a Zeiss LSM800 confocal microscope at the Johns Hopkins University Microscope Facility.

Statistics

Detailed statistical information on data is described in each figure legend. Briefly, no statistical methods were used to predetermine sample sizes, but most of our sample sizes are similar to those generally employed in the field. When performing parametric tests such as *t*-tests, data distribution was assumed to be normal because such parametric tests are known to be robust against the normality assumption. When applying ANOVA of two factors with one within-subjects factor and the other between-subjects factor, two-way repeated measures (RM) ANOVA was applied with the Greenhouse-Geisser correction for the violation of the sphericity assumption. For a within-group comparison, two-tailed paired Student's *t*-test was used.

**Fig. S1. The activity of excitatory neurons in the posterior insular cortex (PIC) is not required for threat memory formation in the trace FC. (A)** Optogenetic inhibition of PIC excitatory neurons during the entire period of each trial did not disrupt threat memory formation in the trace FC paradigm. Entire ($N_{CON}$=12, $N_{Inhibition}$=11; two-way RM ANOVA, Inhibition x CS, $F_{1,21}$=2.048, $p$=0.167). Graphs expressed as mean ± SEM.

### Fig. S1

A

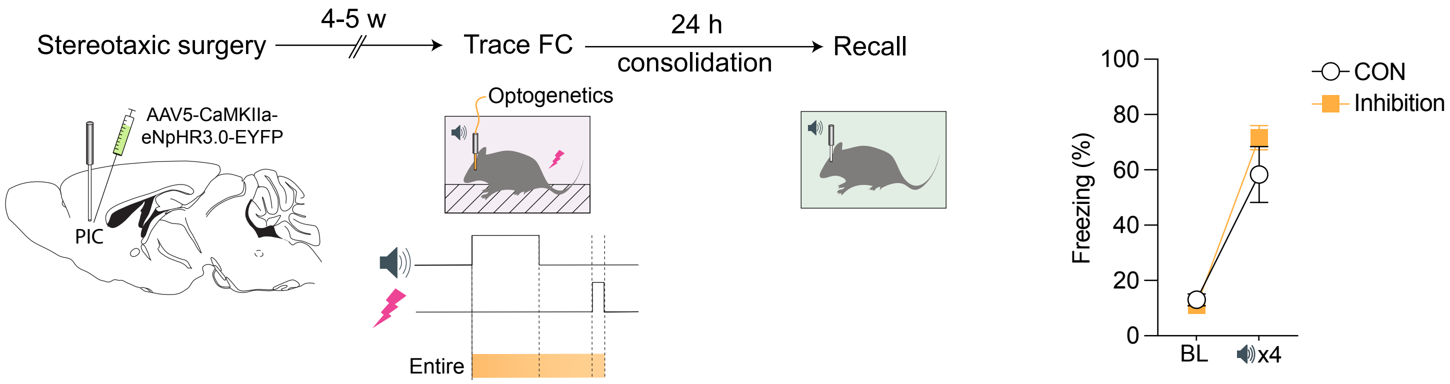

Stereotaxic surgery
4-5 w
Trace FC
24 h
consolidation
Recall
AAV5-CaMKIIa-
eNpHR3.0-EYFP
PIC
Optogenetics
Entire
Freezing (%)
100
80
60
40
20
0
BL
x4
CON
Inhibition
